# A Hierarchical Approach Using Marginal Summary Statistics for Multiple Intermediates in a Mendelian Randomization or Transcriptome Analysis

**DOI:** 10.1101/2020.02.03.924241

**Authors:** Lai Jiang, Shujing Xu, Nicholas Mancuso, Paul J. Newcombe, David V. Conti

## Abstract

**Background:** Previous research has demonstrated the usefulness of hierarchical modeling for incorporating a flexible array of prior information in genetic association studies. When this prior information consists of effect estimates from association analyses of single nucleotide polymorphisms (SNP)-intermediate or SNP-gene expression, a hierarchical model is equivalent to a two-stage instrumental or transcriptome-wide association study (TWAS) analysis, respectively.

**Methods:** We propose to extend our previous approach for the joint analysis of marginal summary statistics (JAM) to incorporate prior information via a hierarchical model (*hJAM*). In this framework, the use of appropriate effect estimates as prior information yields an analysis similar to Mendelian Randomization (MR) and TWAS approaches such as FUSION and S-PrediXcan. However, *hJAM* is applicable to multiple correlated SNPs and multiple correlated intermediates to yield conditional estimates of effect for the intermediate on the outcome, thus providing advantages over alternative approaches.

**Results:** We investigate the performance of *hJAM* in comparison to existing MR approaches (inverse-variance weighted MR and multivariate MR) and existing TWAS approaches (S-PrediXcan) for effect estimation, type-I error and empirical power. We apply *hJAM* to two examples: estimating the conditional effects of body mass index and type 2 diabetes on myocardial infarction and estimating the effects of the expressions of gene *NUCKS1* and *PM20D1* on the risk of prostate cancer.

**Conclusions:** Across numerous causal simulation scenarios, we demonstrate that *hJAM* is unbiased, maintains correct type-I error and has increased power.

**Key Messages:**

- Mendelian randomization and transcriptome-wide association studies (TWAS) can be viewed as similar approaches via a hierarchical model.
- The hierarchal joint analysis of marginal summary statistics (*hJAM*) is a multivariate Mendelian randomization approach which offers a simple way to address the pleiotropy bias that is introduced by genetic variants associated with multiple risk factors or expressions of genes.
- *hJAM* incorporates the linkage disequilibrium structure of the single nucleotide polymorphism (SNPs) in a reference population to account for the correlation between SNPs.
- In addition to Mendelian randomization and TWAS, *hJAM* offers flexibility to incorporate functional or genomic annotation or information from metabolomic studies.

## Introduction

Instrumental variable analysis with genetic variants has been widely used as a general framework for estimating effects of risk factors and gene expression on an outcome (Figure 1)^1–4^. Within this framework using single-nucleotide polymorphisms (SNPs) as instrumental variables, the intermediates *X* can be modifiable risk factors, expression of genes, or other potential intermediates such as methylation, metabolites or proteomics. To be a valid instrumental variable and to yield a causal effect of a risk factor, the genetic variants selected as the instruments must satisfy three assumptions: (1) they must not be associated with the outcome except through the intermediate, (2) they must be at least moderately associated with the intermediate, and (3) they must be independent of potential confounders of the association between the intermediate and the outcome (Figure 1). The violation of the first assumption results in a bias estimate due to pleiotropy. Weak instrument bias will be introduced if the second assumption is violated since the random error may mask the effect of the intermediate on the outcome^5^. Previous work has demonstrated that weak instruments may lead to a large bias in estimators even though the first assumption is only slightly violated^6^. Finally, the law of independent assortment of genetic variants within a homogeneous population or the ability to adequately control for potential confounding due to population structure, often leads genetic variants fulfilling the third assumption.

**Figure 1.**
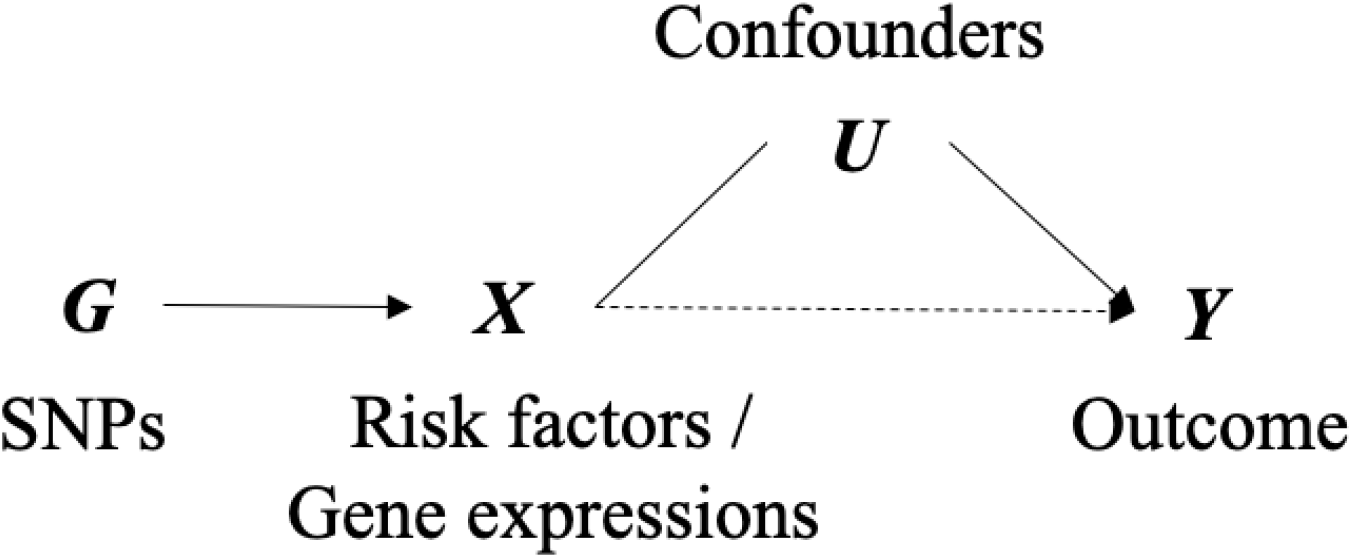
The direct acyclic graph (DAG) for instrumental variable analysis with genetic variants. This DAG describes the framework for several approaches. Arrow denotes a causal effect in the direction of the arrow. Solid line refers to moderate or strong association and dashed line refers to uncertain association.

Mendelian randomization (MR) and transcriptome-wide association studies (TWAS) are the two major approaches within the instrumental variable analysis framework using genetic variants. MR approaches focus on the modifiable risk factors while TWAS approaches adopt gene expression as the intermediate. One advantage of using these tools is the ubiquity of publicly-available genome-wide association studies (GWAS), such as UK Biobank^7^, facilitates researchers to initiate investigation of complex traits and diseases nearly immediately^8^. The existing approaches differ in their strategies to combine the summary data from GWAS or RNA sequencing data.

In this paper, we propose an approach that leverages the joint analysis of marginal summary statistics (*JAM*)^9^, a scalable algorithm designed to analyze published marginal summary statistics from GWAS under a joint multi-SNPs model to identify causal genetic variants for fine mapping. Here, we extend *JAM* with a hierarchical model to incorporate SNP-intermediate association estimates and unify the framework of MR and TWAS approaches when multiple intermediates and/or correlated SNPs exist.

## Methods

### Unify the framework of Mendelian Randomization and TWAS

Instrumental variable analysis with individual-level genotype data can be viewed as a two-stage hierarchical model. Using linear regression, the first stage models the outcome as a function of the genetic variants:

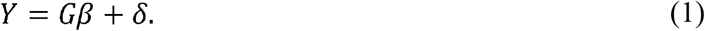

Here, *Y* denotes a n-length vector of a continuous outcome, *G* denotes an *n* × *P* genotype matrix with *P* SNPs and *n* individuals and *δ* denotes the residuals. The second stage models the conditional effect estimates *β* as a function of prior information^10–13^, 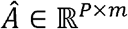:

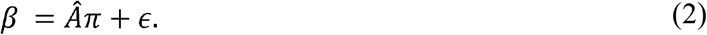

where 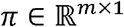 denotes the parameter of interest, the vector of effects for the intermediates *X* on outcome *Y* and *m* is the number of intermediates *X*. We can join these two-stage models into a single linear mixed model by substituting Eq. 2 into Eq. 1^14^:

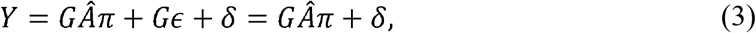

assuming there is no direct effect from the genetic variants to the outcome (i.e. *ϵ* — 0). The estimate of *π* from Eq. 3 is equivalent to the result from the two-stage least square (2SLS) regression, which is employed by PrediXcan^15^ and others^16^. The prior information *Â* is the association estimates between the genetic variants and the intermediate and can be applied to impute the intermediate with the genetic variants:

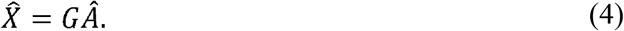

Note that Eq. 4 is the stage-2 in the 2SLS regression and that MR approaches with summary data are developed based on Eq. 2. One key aspect of the instrumental variable analysis with genetic variants is that the *Â* matrix is computed from a separate data, i.e. 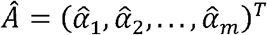, where 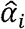 denotes the vector of association estimates between genetic variants and *i^th^* intermediate from external data. Two different 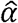 vectors have been used by previous methods. Marginal estimates 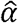 are widely employed by MR where marginal summary statistics from GWAS are being used^17–19^. Conditional estimates of 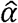, which adjust for the correlations between estimates, can also be incorporated into the framework. One way to construct a conditional estimate 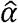 is to apply regularized regression in individual-level data, such as the PredictDB developed for PrediXcan^15^. Another way is to convert the marginal estimates 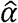 into conditional ones by incorporating the linkage disequilibrium (LD) block among the SNPs using the *JAM* approach^20^. To model multiple intermediates, we construct an *Â* matrix by combining the vectors of effect estimates of the SNPs on each intermediate, 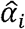, into a matrix with the number of columns equal the number of intermediates (i.e. *m*):

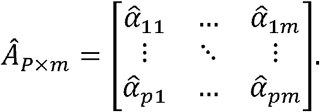

### Hierarchical JAM (hJAM) with summary data

We can employ the same hierarchical model to marginal summary data. Following Newcombe et al.^9^, we use the marginal summary statistics, 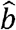, which are obtained from a GWAS and the minor allele frequency (MAF) of the genetic variants, 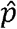, to construct a vector *z* representing the genotype weighted effect for each genetic variant *i*:

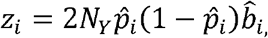

assuming Hardy-Weinberg Equilibrium. The MAF can be extracted from the same GWAS or using external populations such as 1000 Genomes Project^21^ as reference data. Using standard linear algebra, we can express the distribution of *z* as

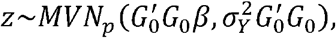

where 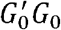 denotes the *P* × *P* genotype variance-covariance from a centered reference data set (e.g. 1000 Genome) to obtain the conditional effects of SNPs on the outcome, *β*. Details are described in Newcombe et al.^9^. To simplify the likelihood, we perform a Cholesky decomposition transformation 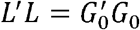. Then, we transform *z* into *z_L_* with the inverse of *L*′ as *z_L_* = *L*′^−1^*z*. When *L* is positive semi-definite, we add a ridge term, i.e. a small positive element, on the diagonal to enforce it to be a positive definite matrix. The regularization term has a very small effect on the estimates while guaranteeing the invertibility of the *L* matrix. Then, the *z_L_* is a vector of independent statistics that can be expressed as

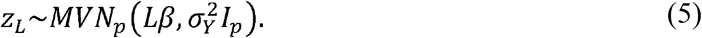

Similar to above, we then fit a hierarchical model by incorporating the second-stage model (Eq.2) into Eq.5 and construct the *hJAM* model as

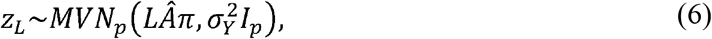

assuming no direct effect from genetic variants to the outcome. Here, 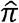 denotes the association parameter of interest between the intermediate and outcome and is estimated using maximum likelihood and the statistical significance is given by a Wald test. The estimate of 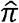 and corresponding variance are

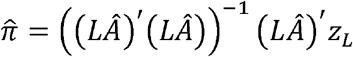

and

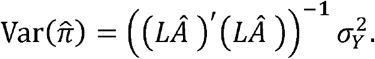

Egger-type approaches can be implemented in this framework by allowing an intercept in Eq. 6 by adding a column of ones to *Â* matrix, which is analogue to MR Egger regression.

#### Simulation studies

To assess the performance of *hJAM*, we performed an extensive set of simulation studies. For each simulation, we simulated three standardized individual genotype matrices *G_X_*, *G_Y_*, and *G_L_*, an intermediate matrix *X*, and an outcome vector *Y*. We then generated the summary statistics, including marginal effects 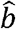, *Â*, and the reference LD structure, from the individual-level data.

For each genotype matrix, we had two inter-block relationships: no LD and moderate LD (*R* – 0.6). Each SNP block (i.e. *G*_1_,, *G*_2_ and *G*_3_ in Figure 2) contains 10 SNPs, in which we set 3 SNPs to be causal to the intermediate with 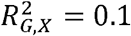. The MAF was sampled from a uniform distribution (0.05, 0.3). Sample size for each genotype data set was set to be *N_G_X__* = 1000, *N_G_Y__* = 5000, and *N_G_L__* = 500, respectively. We simulated two *X*’s and four scenarios representing different causal models for the two intermediates likely to be encountered in epidemiologic studies (Figure 2). For scenario A, *X*_1_ and *X*_2_ were independent. For scenarios B and D, Apnd *X*_2_ were correlated through a shared SNPs set *G3*. The coefficient *λ* in the causal scenarios (Figure 2 (C) and (D)) was simulated by 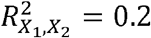. These simulation scenarios are similar to those described in Sanderson et al.^23^.

**Figure 2.**
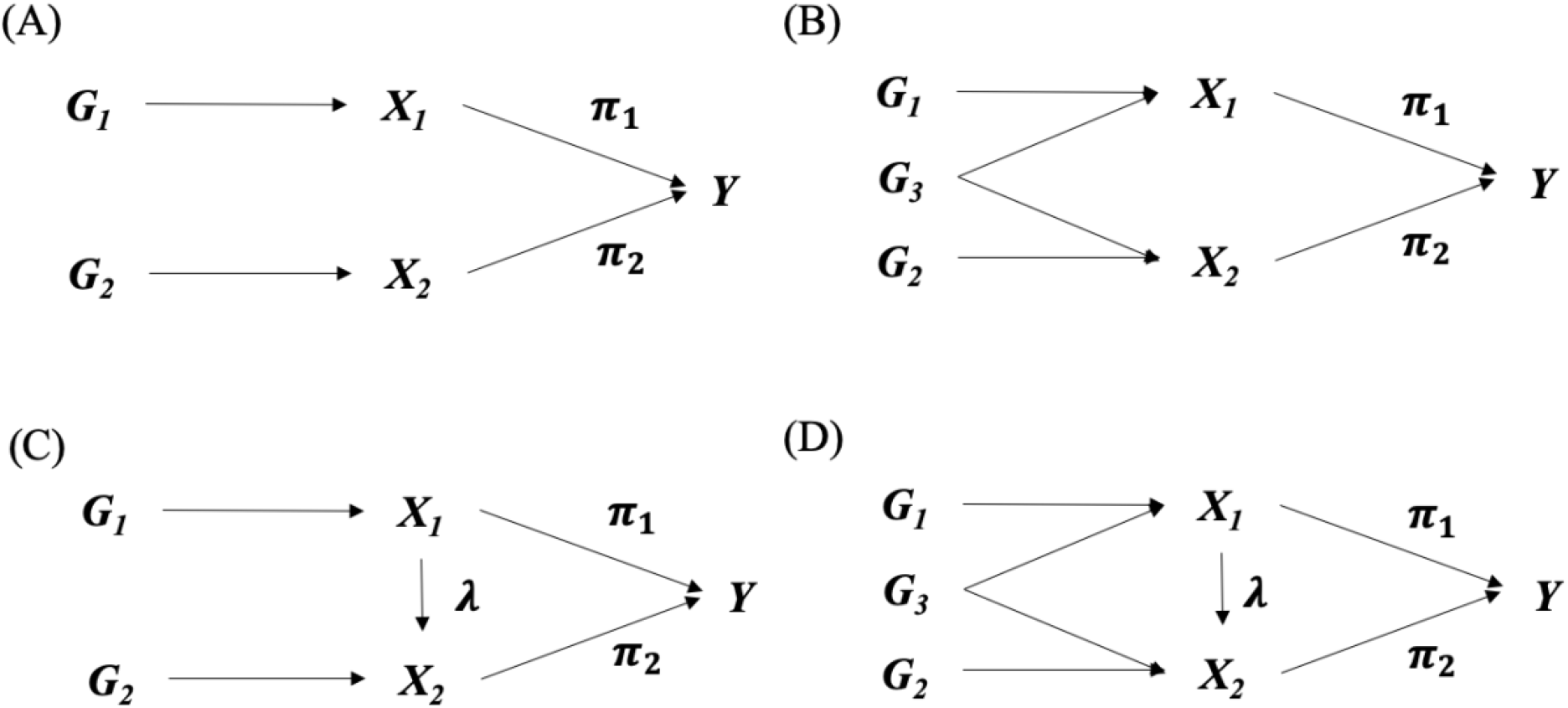
Simulation scenarios of different relationships between X’s. (A) X_1_ and X_2_ are independent. (B) X_1_ and X_2_ are correlated. (C) X_1_ causes X_2_. (D) X_1_ causes X_2_ and correlated.

The primary objective was to estimate 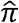 with each true *π_i_* being set to null (*π_i_* = 0) or a positive effect (π_*i*_ = 0.1). To mimic applied applications and to ensure selection of at least two or more SNPs, a forward selection on *Â* was performed to exclude the noninformative variants with a threshold *P* < 0.2 in the analysis step. We compared the performance of our approach to inverse-variance weighted MR (IVW MR)^17^, multivariate inverse-variance weighted MR (MVIVW MR)^18^, and S-PrediXcan^24^ (see Appendix). All simulation analyses were performed in R version 3.4.0. Results were calculated from 1000 replications for each scenario. All tests were two-sided with a type-I error of 0.05.

### Simulation results

The average estimate and standard error for *π* across the four simulation scenarios are presented in Figure 3 and supplementary Table 1 to Table 4, respectively. Supplementary Figure 1 (independent SNPs) and Figure 4 (correlated SNPs) present results for empirical power.

**Figure 3.**
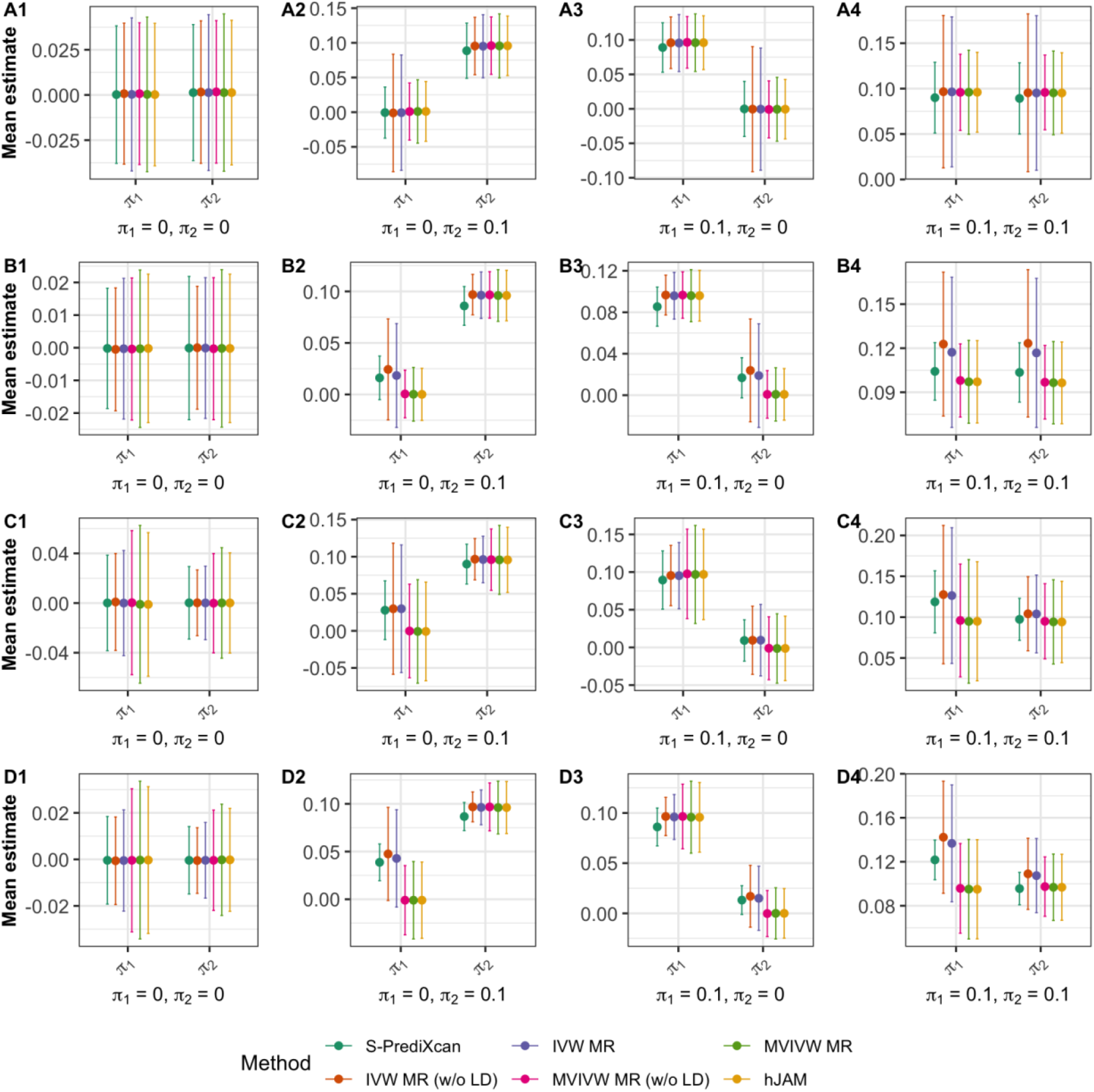
Average estimates and 95% confidence intervals of the correlated SNPs scenarios across 1000 replications. (A) X_1_ and X_2_ are independent. (B) X_1_ and X_2_ are correlated. (C) X_1_ causes X_2_. (D) X_1_ causes X_2_ and correlated. The black solid line refers to the default Type-I error, α – 0.05.

**Figure 4.**
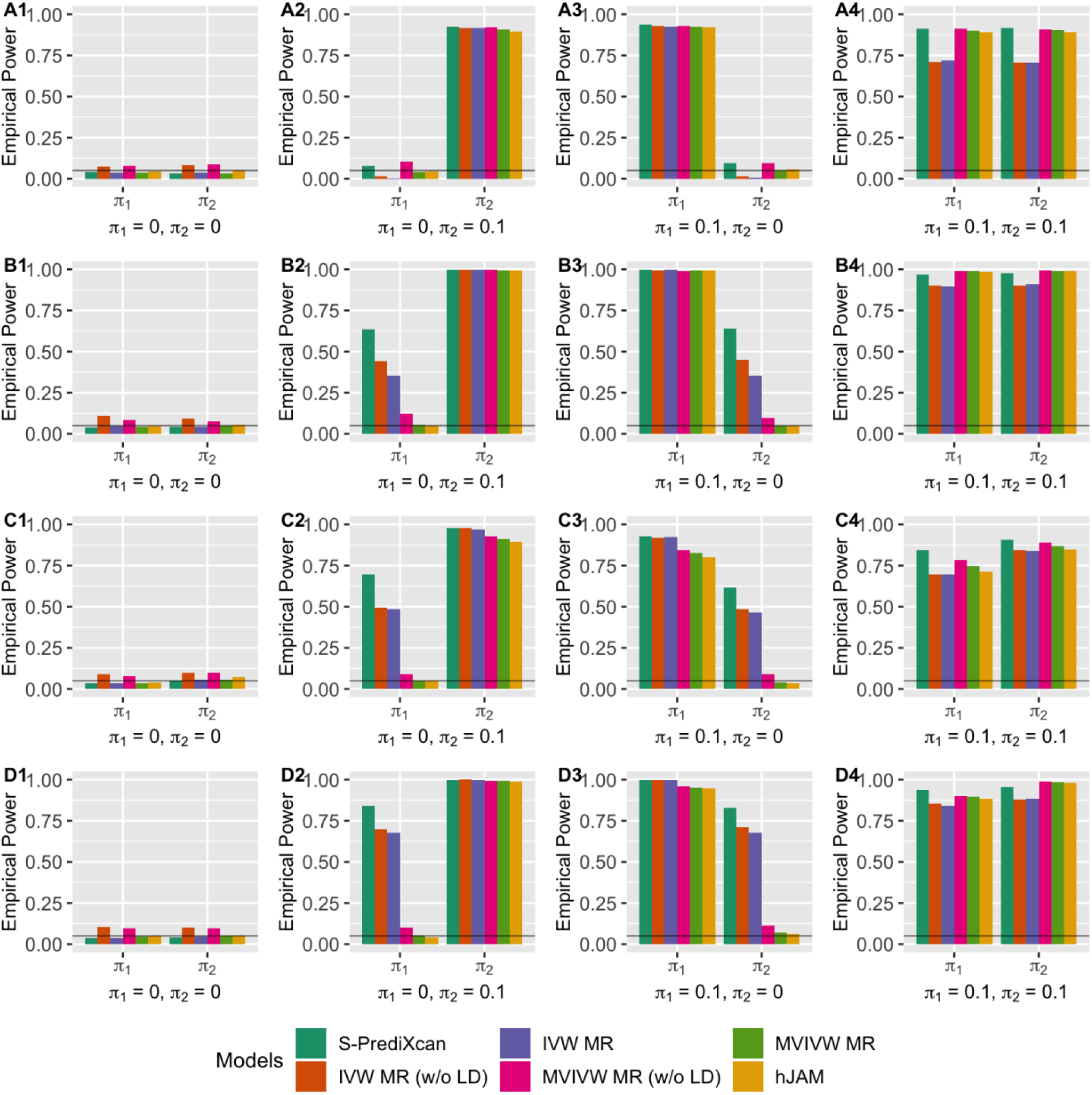
Empirical Power of the correlated SNPs scenarios across 1000 replications. (A) X_1_ and X_2_ are independent. (B) X_1_ and X_2_ are correlated. (C) X_1_ causes X_2_. (D) X_1_ causes X_2_ and correlated. The black solid line refers to the default Type-I error, α – 0.05.

Results from the base scenario A, where *X*_1_ and *X*_2_ were independent, demonstrate that the estimates from most methods were unbiased. However, when IVW MR and MVIVW MR do not incorporate the LD structure, there is a slightly inflated type-I error under simulation scenarios with correlated SNPs. IVW MR with and without correlation had a less precise estimate and lower power compared to the other methods in scenario A (Table 1). When a pleiotropic effect was simulated for each intermediate (scenario B to D), the estimates from *hJAM* and MVIVW MR with LD were unbiased and had a correct type-I error for the corresponding intermediate (Figure 4). The estimates from MVIVW MR without LD were unbiased but showed an inflated type-I error due to a smaller estimated standard error in scenarios in which SNPs were correlated (Figure 4). IVW MR and S-PrediXcan had a biased estimate and an inflated type-I error regardless of the correlation structure of the SNPs. The results for MVIVW MR and IVW MR reflect specification of the LD structure for the instruments when using the *MedelianRandomization*^25^ package. Results without the LD structure showed a poor performance as indicated by increased type-I errors.

**Table 1.** Causal odds ratios (95% confidence interval) for myocardial infarction per unit in body mass index and having type 2 diabetes.

| Methods | Odds ratios (95% CI) | P |
| --- | --- | --- |
| **BMI** |  |  |
| hJAM | 1.38 (1.22, 1.56) | 3.19E-07 |
| MVIVW MR | 1.37 (1.22, 1.54) | 1.94E-07 |
| MVIVW MR (w/o LD) | 1.34 (1.20, 1.49) | 1.65E-07 |
| IVW MR | 1.54 (1.32, 1.79) | 2.07E-08 |
| IVW MR (w/o LD) | 1.53 (1.32, 1.77) | 1.45E-08 |
| S-PrediXcan | 1.66 (1.58, 1.74) | 9.88E-96 |
| Egger-intercept | 0.005 (-0.003, 0.013) | 2.00E-01 |
| **T2D** |  |  |
| hJAM | 1.16 (1.12, 1.20) | 4.12E-11 |
| MVIVW MR | 1.15 (1.11, 1.19) | 8.34E-12 |
| MVIVW MR (w/o LD) | 1.16 (1.11, 1.20) | 1.29E-11 |
| IVW MR | 1.15 (1.11, 1.20) | 1.77E-14 |
| IVW MR (w/o LD) | 1.15 (1.11, 1.20) | 1.98E-14 |
| S-PrediXcan | 1.14 (1.11, 1.16) | 9.43E-109 |
| Egger-intercept | 0.007 (0.000, 0.013) | 0.017 |

Abbreviation: w/o LD, without linkage disequilibrium adjustment; s.e., standard error; 95% CI, 95% confidence interval.
Note: \* For MR Egger (intercept), we showed log odds ratio and its 95% CI.

---

**Supplementary Table 1 The estimate and its standard error of simulation scenario A: independent *X*’s.**

**Supplementary Table 2 The estimate and its standard error of simulation scenario B: correlated *X*’s.**

**Supplementary Table 3 The estimate and its standard error of simulation scenario C: *X*_1_ causes X_2_.**

**Supplementary Table 4 The estimate and its standard error of simulation scenario D: *X*_1_ causes *X*_2_ and *X*_1_ and *X*_2_ are correlated**.

**Supplementary Figure 1 Empirical Power of the independent SNPs scenarios across 1000 replications.**

(A) X_1_ and X_2_ are independent. (B) X_1_ and X_1_ are correlated. (C) X_1_ causes X_1_. (D) X_1_. causes X_2_ and correlated. The black solid line refers to the default Type-I error, α – 0.05

## Data Examples

To demonstrate *hJAM* on real data, we applied various methods to two examples: 1) for body mass index (BMI) and type 2 diabetes (T2D) on myocardial infarction (MI); and 2) gene expression and prostate cancer risk. As the study populations for both examples include individuals of European ancestry, we used the 503 European-ancestry subjects from the 1000 Genomes Project^21^ as our reference data for the LD structure.

### Causal effect of BMI and T2D on myocardial infarction

Previous studies have shown that obesity^26,27^ and T2D^28, 29^ are two important risk factors for MI. In addition, the association between obesity and T2D is well-established^30, 31^. A directed acyclic graph (DAG) shows the relationships between the two risk factors and MI (Figure 5).

**Figure 5.**
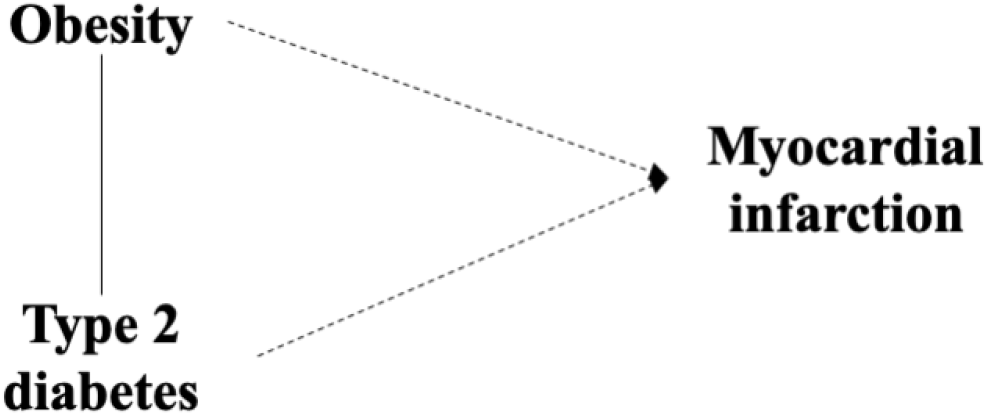
Direct acyclic graph (DAG) of the relationship between BMI, type 2 diabetes and myocardial infarction.

To examine the conditional effects of the two risk factors, we extracted the summary statistics for MI, BMI, and T2D from the UK Biobank (n = 459,324)^7^, GIANT consortium (n = 339,224)^32^, and DIAGRAM+GERA+UKB (n = 659,316)^33^, respectively. In total, 75 SNPs and 136 SNPs were identified as genome-wide significant for BMI and T2D, respectively (supplementary Figure 2 and supplementary Figure 3). In this set of SNPs, there was one overlapping SNP in both the instrument sets for BMI and T2D (rs7903146, *α*_BMI_ = −0.016, *P*_BMI_ = 1.4 × 10^−14^, *α*_T2D_ = 0.319, and *P*_T2D_ = 1.7 × 10^−204^). This SNP is a well-known T2D associated SNP and has being identified as a BMI-associated hit in GIANT. Additionally, four correlated pairs of SNPs exist between the two sets (supplementary Table 5). We re-orientated the effects of all SNPs but one (except the effect of the overlapping SNP rs7903146 on BMI) to have a positive effect and we used MR Egger regression^22^ and *hJAM* Egger to detect a potential directional pleiotropy effect.

Results are shown in Table 1. All methods suggested a significantly increasing risk of MI with an increased BMI and the presence of T2D. This agrees with previous studies^26, 28^. The magnitude of *hJAM* and MVIVW MR were similar while IVW MR and S-PrediXcan showed larger estimated values. The odds ratio (OR) from *hJAM* for the risk of MI was 1.38 (95% CI=1.22, 1.56) and 1.16 (95% CI=1.12, 1.20) for per one unit increase in BMI and having T2D, respectively. MVIVW MR with LD has similar estimates with 1.37 (95% CI=1.22, 1.54) and 1.15 (95% CI=1.11, 1.19) for BMI and having T2D, respectively. The difference in estimates between the multivariate approaches and the univariate MR/TWAS approaches may be attributed to potential pleiotropy not accounted for in the analyses that do not model the intermediates jointly. When modeled jointly, results from *hJAM* Egger suggested that there was no residual pleiotropy detected when we incorporated both BMI- and T2D-associated instruments in the analysis (*P* = 0.57). In contrast, the MR-Egger approach applied univariately to T2D resulted in a significant test for the intercept, suggesting the presence of pleiotropy, potentially due to association of some of the SNPs to the outcome via BMI.

**Supplementary Table 5 Four correlated pairs of SNPs in the instrument sets of BMI and type 2 diabetes.**

**Supplementary Figure 2 Scatter plots for the univariate effect estimates 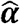 vs. 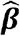 for BMI (A) and type 2 diabetes (B).**

**Supplementary Figure 3 Heatmap of the Pearson correlation between the 210 instrumental SNPs in data example 1: BMI and type 2 diabetes of myocardial infarction.**

### Causal effect of PM20D1 and NUCKS1 on prostate cancer risk

To further illustrate the benefit of *hJAM*, we next considered the gene-prostate cancer risk association of two genes on chromosome 1q32.1, gene *PM20D1* (Peptidase M20 Domain Containing 1) and gene *NUCKS1* (Nuclear Casein Kinase and Cyclin Dependent Kinase Substrate 1). Both *PM20D1* and *NUCKS1* are protein coding genes and previous transcriptome studies have found a significant effect of both *PM20D1* and *NUCKS1* on the risk of prostate cancer among a European-ancestry population^34, 35^. Due to the close proximity of the two genes along the genome, there is a potential for a univariate approach to result in biased estimates. To examine the effects jointly, we applied *hJAM* to this research question.

We constructed the *Â* matrix with 114 marginally significant expression quantitative trait loci (eQTL) estimates for the two genes from GTEx v7^36^. Among the 114 eQTLs, one eQTL has significant associations with both *PM20D1* and *NUCKS1*. To limit the correlation between the eQTLs (|*R*| > 0.9), we used priority pruner^37^ to prune the eQTLs by limited the absolute pairwise correlation coefficient |*R*| < 0.7 and using the magnitude of the eQTLs association for each gene as the priority criteria. After pruning, we had 12 eQTLs in the analysis set (supplementary Figure 3). The genome-wide summary statistics for the risk of prostate cancer was taken from a published GWAS with more than 140,000 European-ancestry men^38^.

Table 2 presents results from *hJAM* and the competing approaches. *hJAM* and MVIVW MR with LD yield non-significant results for both *PM20D1* and *NUCKS1* for the risk of prostate cancer (*P*_PM20D1_ = 0.90 and *P*_NUCKS1_ = 0.21 for hJAM, and *P*_PM20D1_ = 0.09 and *P*_NUCKS1_ – 0.17 for MVIVW MR with LD). However, univariate models, including IVW MR and S-PrediXcan, results in a significant positive effect for prostate cancer risk for *PM20D1* and *NUCKS1* (*P*_PM20D1_ = 0.024 and *P*_NUCKS1_ = 3.53 × 10^−15^ for IVW MR without correlation, and *P*_PM20D1_ – 0.003 and *P*_NUCKS1_ = 2.84 × 10^−10^ for S-PrediXcan). We consider the significance in the univariate models was due to the correlation between the two genes and the LD between the eQTLs, which could be adjusted for by the *hJAM* and MVIVW MR with LD models.

**Table 2.** Causal odds ratios (95% confidence interval) for prostate cancer risk per unit increasing in gene expression reads.

| Methods | Odds ratio (95% CI) | P |
| --- | --- | --- |
| ***PM20D1*** |  |  |
| hJAM | 0.10 (0.92, 1.08) | 0.91 |
| MVIVW MR | 0.10 (0.93, 1.07) | 0.90 |
| MVIVW MR (w/o LD) | 0.99 (0.94, 1.04) | 0.66 |
| IVW MR | 1.02 (0.96, 1.10) | 0.49 |
| IVW MR (w/o LD) | 1.03 (1.00, 1.05) | 0.02 |
| S-PrediXcan | 1.01 (1.00, 1.01) | 0.003 |
| ***NUCKS1*** |  |  |
| hJAM | 1.12 (0.93, 1.36) | 0.21 |
| MVIVW MR | 1.12 (0.95, 1.33) | 0.17 |
| MVIVW MR (w/o LD) | 1.15 (0.10, 1.33) | 0.06 |
| IVW MR | 1.16 (1.10, 1.21) | 5.03E-10 |
| IVW MR (w/o LD) | 1.16 (1.12, 1.20) | 3.53E-15 |
| S-PrediXcan | 1.10 (1.07, 1.13) | 2.84E-10 |

Abbreviation: w/o LD, without linkage disequilibrium adjustment; s.e., standard error; 95% CI, 95% confidence interval.

**Supplementary Figure 4 Heatmap of the Pearson correlation between the 12 instrumental SNPs in data example 2: the effect of two gene expressions on the risk of prostate cancer.**

## Discussion

In this paper, we have proposed a two-stage hierarchical model which unifies the framework of Mendelian randomization and transcriptome-wide association tools and can be applied to correlated instruments and multiple intermediates. We have implemented the method in an R package which is available on Github (https://github.com/lailylajiang/hJAM).

When only one intermediate or multiple independent intermediates present, *hJAM* yield an equivalent estimate and standard error to alternative approaches (see Appendix). However, when intermediates are correlated, only MVIVW MR showed a comparable performance with *hJAM* under the independent SNPs scenarios. For correlated SNPs scenarios, when the LD structure is specified, *hJAM’s* estimates are empirically equivalent to MVIVW MR although the two approaches use slightly different weighted matrices – *hJAM* uses the adjusted variance-covariance matrix of SNPs from a reference panel while MVIVW MR uses an inverse-variance matrix. Nevertheless, we believe that the *hJAM* formulation offers several advantages in flexibility to specify the *Â* matrix. As in TWAS, this matrix can specify eQTL estimates or as in more classical MR approaches this can specify SNP-intermediate associations. Moreover, it can incorporate other types of prior information such as functional or genomic annotation or information from metabolomic studies. Inclusion of this type of annotation information can offer potential advantages for characterization of SNP effects as demonstrated in the hierarchical modeling context^10, 11, 40^. Future research needs to be performed on how best to construct this matrix for various types of intermediates.

Although *hJAM* provides an overall improvement over most existing MR methods, it is also susceptible to the caveats of these types of approaches. Firstly, it may subject to the bias in estimation due to unknown horizontal pleiotropy. *hJAM* can be extended to include a column of ones to the *Â* matrix to allow for estimating an intercept term to formally test for directional pleiotropy, analogous to MR Egger^22^. This *hJAM-Egger* version showed a similar performance to the univariate MR-Egger regression with unbiased estimates under simulations in which the horizontal pleiotropy is balanced, but bias estimates in the presence of unbalanced pleiotropy (results not shown)^22^. *hJAM-Egger* can be applied as a sensitivity analysis of a multivariable framework MR analysis^41^. An extension of the current *hJAM* approach could include variable selection to assess the pleiotropy assumption before incorporating the *Â* matrix into the model. Several approaches have been proposed, such as JAM MR^42^ and MR-presso^43^. Secondly, the effects of the SNPs on the intermediates and the outcome, and the causal effect of intermediates on the outcome may be non-linear (e.g. interactions). One way to address such limitation is to use summary data from stratified GWAS; however, it may attenuate the power due to a smaller sample size of the subset GWAS.

In applied applications, population structure may introduce potential difficulties for *hJAM*, as is similar for all MR and TWAS approaches using summary statistics. First, there is the reliance that the association statistics are unbiased due to potential confounding by population structure. This includes summary data for the SNPs to intermediate associations in *Â* matrix, as well as the marginal SNP-outcome associations using within the *hJAM* model. However, given that modern techniques to account for population structure are often sufficient^44, 45^, this is a fair assumption. Additionally, to account for the correlation structure between SNPs, *hJAM* assumes that the LD structure estimated from the reference data is the same as the study data used to generate the summary statistics. Since *hJAM* and MVIVW MR incorporate the correlation structure of SNPs in a slightly different weight matrices, there is the potential for this to impact these methods differently. Although, in a limited set of simulations we found that both methods are fairly robust to scenarios in which the reference data and the association data have modest differences in LD structures (results not shown).

In contrast to most current methods that rely on independent SNPs or analyze intermediates in isolation, we propose a two-stage hierarchical model to jointly model summary statistics (*hJAM*) for correlated SNPs and multiple intermediates within Mendelian Randomization and transcriptome-wide association studies. As technology expands the potential use of these types of studies to proteomic, methylation and metabolomic data, such flexible approaches will be needed to account for the potential increase in complexity in underlying relationships between factors.

## Acknowledgement

The authors thank Drs. Duncan C. Thomas, William Gauderman, and Juan Pablo Lewinger for valuable discussions and comments throughout development.

