## Appendix for "A Hierarchical Approach Using Marginal Summary Statistics for Multiple Intermediates in a Mendelian Randomization or Transcriptome Analysis"

We compared the performance of our approach to inverse-variance weighted MR (IVW MR)^1^, multivariate inverse-variance weighted MR (MVIVW MR)^2^, and S-PrediXcan^3^. The default versions of IVW MR^1^ and MVIVW MR^2, 4^ do not specify a correlation structure between SNPs. However, the R package for implementation, *MendelianRandomization*^5^, provides functionality to incorporate the LD structure in the analysis. Thus, for IVW MR and MVIVW MR, we implemented them in two ways: with or without specifying the LD structure of the simulated SNPs. Inverse-variance weighted MR^1^ (IVW MR) is a classic MR method which applies a fixed-effect meta-analysis approach to combine the ratio estimates using each variate from two GWAS. The point estimate from IVW MR is similar in spirit to the second-stage model of the hierarchical model and is equivalent to a weighted linear regression without an intercept of SNP-outcome (*Y* and *G*) associations on SNP- intermediate ($X_{i}$ and *G*) associations. MVIVW MR is an extension on IVW MR which allows multiple intermediates and uses multivariate weighted linear model to test the joint effects of multiple intermediates^5^. S-PrediXcan is an extension of PrediXcan using summary statistics as an input^3^ to estimate the univariate gene expression-outcome associations.

**Hierarchical Joint Analysis of Summary Statistics (*hJAM*)**

To theoretically compare with other approaches, we write *hJAM* estimates in another form under several assumptions. Use the distribution as shown in Eq. 6 and assume that the SNPs are *independent*, we can simplify the estimate of *hJAM* as

$$\hat{\pi}_{\mathrm{hJAM}}=\left( \left( \boldsymbol{L}\hat{\boldsymbol{A}} \right)^{\boldsymbol{'}}\left( \boldsymbol{L}\hat{\boldsymbol{A}} \right) \right)^{\boldsymbol{-1}}\left( \boldsymbol{L}\hat{\boldsymbol{A}} \right)\boldsymbol{'z}_{\boldsymbol{L}}=\left( {\hat{\boldsymbol{A}}}^{\boldsymbol{'}}\boldsymbol{L}^{\boldsymbol{'}}\boldsymbol{L}\hat{\boldsymbol{A}} \right)^{\boldsymbol{-1}}{\hat{\boldsymbol{A}}}^{\boldsymbol{'}}\boldsymbol{L}^{\boldsymbol{'}}{\boldsymbol{L}^{\boldsymbol{'}}}^{\boldsymbol{-1}}\boldsymbol{z}=\left( {\hat{\boldsymbol{A}}}^{\boldsymbol{'}}\boldsymbol{G}_{\boldsymbol{0}}^{\boldsymbol{'}}\boldsymbol{G}_{\boldsymbol{0}} \hat{\boldsymbol{A}} \right)^{\boldsymbol{-1}}{\hat{\boldsymbol{A}}}^{\boldsymbol{'}}\boldsymbol{G}_{\boldsymbol{0}}^{\boldsymbol{'}}\boldsymbol{y}=\frac{1}{N_{Y}}\left( {\hat{\boldsymbol{A}}}^{\boldsymbol{'}}\boldsymbol{\Gamma}_{\boldsymbol{g}} \hat{\boldsymbol{A}} \right)^{\boldsymbol{-1}}{\hat{\boldsymbol{A}}}^{\boldsymbol{'}}\boldsymbol{G}_{\boldsymbol{0}}^{\boldsymbol{'}}\boldsymbol{G}_{\boldsymbol{0}} \hat{\boldsymbol{\beta}}\boldsymbol{=}\left( \hat{\boldsymbol{A}} \boldsymbol{\Gamma}_{\boldsymbol{g}} \hat{\boldsymbol{A}} \right)^{\boldsymbol{-1}}{\hat{\boldsymbol{A}}}^{\boldsymbol{'}}\boldsymbol{\Gamma}_{\boldsymbol{g}} \hat{\boldsymbol{\beta}}\boldsymbol{,}$$

with variance being estimated as

$$\text{Var}\left( {\hat{\boldsymbol{\pi}}}_{\mathrm{hJAM}} \right)\boldsymbol{=}\left( \left( \boldsymbol{L}\hat{\boldsymbol{A}} \right)^{\boldsymbol{'}}\left( \boldsymbol{L}\hat{\boldsymbol{A}} \right) \right)^{\boldsymbol{-1}}\sigma_{Y}^{2}={\frac{1}{N_{Y}}\left( {\hat{\boldsymbol{A}}}^{\boldsymbol{'}}\boldsymbol{\Gamma}_{\boldsymbol{g}}\hat{\boldsymbol{A}} \right)}^{\boldsymbol{-1}}\sigma_{Y}^{2},$$

Then, denote $\sigma_{g}^{\boldsymbol{2}}\boldsymbol{=}{\hat{\boldsymbol{A}}}^{\boldsymbol{'}}\boldsymbol{\Gamma}_{\boldsymbol{g}} \hat{\boldsymbol{A}}$, when $M=1$ with all instruments being independent, we have

$$\hat{\pi}_{\mathrm{hJAM}}=\frac{1}{N_{Y}\hat{\sigma}_{g}^{\boldsymbol{2}}}{\hat{\boldsymbol{A}}}^{\boldsymbol{'}}\boldsymbol{G}_{\boldsymbol{0}}^{\boldsymbol{'}}\boldsymbol{G}_{\boldsymbol{0}} \hat{\boldsymbol{\beta}} =\frac{1}{N_{Y}\hat{\sigma}_{g}^{\boldsymbol{2}}} \sum_{k} \hat{\alpha}_{k}\cdot\hat{\beta}_{k}{\cdot g}_{kk}^{2}=\frac{\sum_{k} \hat{\alpha}_{k}\hat{\beta}_{k} \hat{\sigma}_{g, k}^{2}}{\sum_{k} \hat{\alpha}_{k}^{2} \hat{\sigma}_{g, k}^{2}},$$

where $g_{kk}^{2}$ denotes the $\left( k, k \right)^{th}$ element of $\boldsymbol{G}_{\mathbf{0}}^{\mathbf{'}}\boldsymbol{G}_{\mathbf{0}}$**,** $\hat{\sigma}_{g, k}^{2}=\frac{g_{kk}^{2}}{N_{Y}}$ denotes the variance of the SNP k and  $\hat{\sigma}_{g}^{2}= {\hat{\boldsymbol{A}}}^{\mathbf{'}}\boldsymbol{\Gamma}_{\boldsymbol{g}} \hat{\boldsymbol{A}} \mathbf{=}\sum_{k} \alpha_{k}\cdot\hat{\sigma}_{g,k}^{2}\cdot\alpha_{k}= \sum_{k} \alpha_{k}^{2} \hat{\sigma}_{g,k}^{2}$.

For the variance of the estimate, we have

$$\text{Var(}\hat{\pi}_{\mathrm{hJAM}})={\frac{\hat{\sigma}_{Y}^{2}}{N_{Y}}\left( {\hat{\boldsymbol{A}}}^{\boldsymbol{'}}\boldsymbol{\Gamma}_{\boldsymbol{g}}\hat{\boldsymbol{A}} \right)}^{\boldsymbol{-1}}=\frac{\hat{\sigma}_{Y}^{2}}{N_{Y}\hat{\sigma}_{g}^{\boldsymbol{2}}}.$$

The Z test statistic is

$$Z_{\text{hJAM}}=\frac{\hat{\pi}_{hJAM}}{\sqrt{\text{Var(}\hat{\pi}_{hJAM})}}=\frac{\sum_{k} \hat{\alpha}_{k}\hat{\beta}_{k} \hat{\sigma}_{g, k}^{2}}{\hat{\sigma}_{g}^{2}}\cdot\sqrt{\frac{N_{Y}\hat{\sigma}_{g}^{\boldsymbol{2}}}{\hat{\sigma}_{Y}^{2}}}=\frac{\sum_{k} \hat{\alpha}_{k}\hat{\beta}_{k} \hat{\sigma}_{g, k}^{2}}{\hat{\sigma}_{g}}\cdot\sqrt{\frac{N_{Y}}{\hat{\sigma}_{Y}^{2}}}.$$

#### Multivariate Inverse-variance Weighted Mendelian Randomization

We followed the function of MVIVW MR in R package *MendelianRandomization* ^5^ for theoretical comparison. When no LD specified, the solutions of the estimates and corresponding standard errors are the weighted least square maximum likelihood estimate solutions of the regression

$$\hat{\boldsymbol{\beta}}\boldsymbol{\sim}\hat{\boldsymbol{A}}\boldsymbol{,}$$

and the solutions are

$${\hat{\boldsymbol{\pi}}}_{\mathrm{MVIVW}}=\left( {\hat{\boldsymbol{A}}}^{\boldsymbol{'}}\boldsymbol{W}_{\mathrm{MVIVW}}\hat{\boldsymbol{A}} \right)^{-1} {\hat{\boldsymbol{A}}}^{\boldsymbol{'}}\boldsymbol{W}_{\mathrm{MVIVW}} \hat{\boldsymbol{\beta}}$$

and

$$\text{Var}\left( {\hat{\boldsymbol{\pi}}}_{\mathrm{MVIVW}} \right)= \text{diag}\left( \left( {\hat{\boldsymbol{A}}}^{\boldsymbol{'}} \boldsymbol{W}_{\mathrm{MVIVW}} \hat{\boldsymbol{A}} \right)^{\boldsymbol{-1}} \right)\boldsymbol{,}$$

where $\boldsymbol{W}_{\text{MVIVW,independent}}$ is a diagonal matrix with the inverse standard errors of  $\hat{\beta}$, i.e.

$$\boldsymbol{W}_{\text{MVIVW,independent}}= \left( \begin{matrix} \frac{1}{\text{se}\left( \hat{\beta}_{1} \right)^{2}} & \cdots& 0 \\ \vdots& \ddots& \vdots\\ 0 & \cdots& \frac{1}{\text{se}\left( \hat{\beta}_{p} \right)^{2}} \end{matrix} \right)=\frac{N_{Y}}{\hat{\sigma}_{Y}^{2}}\cdot\left( \begin{matrix} \hat{\sigma}_{g,1}^{2} & \cdots& 0 \\ \vdots& \ddots& \vdots\\ 0 & \cdots& \hat{\sigma}_{g,p}^{2} \end{matrix} \right).$$

Here, $N_{Y}$ denotes the sample size of $G_{Y}$. In *hJAM*, we have a similar form of the estimate as MVIVW MR. If we rewrite the variance of the estimates from hJAM as $\mathrm{Var}\left( \hat{\pi}_{\mathrm{hJAM}} \right)=\mathrm{diag}\left( \left( {\hat{\boldsymbol{A}}}^{\boldsymbol{'}} \boldsymbol{W}_{\mathrm{hJAM}} \hat{\boldsymbol{A}} \right)^{\boldsymbol{-1}} \right)$, we can express the weight matrix of hJAM as

$$\boldsymbol{W}_{\text{hJAM}}=\frac{N_{Y}}{\hat{\sigma}_{Y}^{2}}\cdot\boldsymbol{\Gamma}_{\boldsymbol{g}}=\frac{N_{Y}}{\hat{\sigma}_{Y}^{2}}\cdot\left( \begin{matrix} \hat{\sigma}_{g,1}^{2} & \cdots& \hat{\sigma}_{g,1}\cdot\hat{\sigma}_{g, p}\cdot\rho_{1p} \\ \vdots& \ddots& \vdots\\ \hat{\sigma}_{g,p}\cdot\hat{\sigma}_{g, 1}\cdot\rho_{p1} & \cdots& \hat{\sigma}_{g,p}^{2} \end{matrix} \right),$$

where $\rho_{ij}$ denotes the correlation coefficient between $i^{th}$ SNP and $j^{th}$ SNP, obtaining from the reference panel (i.e. $G_{L}$). Our estimate is equivalent to the estimate from MVIVW MR when SNP are independent, i.e. $\rho_{ij}=0$ for $\forall i\neq j$. Note that, if the sample sizes of the reference panel, $\boldsymbol{G}_{\boldsymbol{L}}$, and that of $\boldsymbol{G}_{\boldsymbol{Y}}$ are different, we will modify the $\boldsymbol{G}_{\boldsymbol{0}}^{\boldsymbol{'}}\boldsymbol{G}_{\boldsymbol{0}}$ matrix with the summary statistics from $\boldsymbol{G}_{\boldsymbol{Y}}$ and scaling the variance and co-variances accordingly (details in the supplementary materials of ^6^). When correlation specified, the weight matrix $\boldsymbol{W}$ of MVIVW MR changes into

$$\boldsymbol{W}_{\text{MVIVW, correlated}}=\left( \left( \mathbf{B}\otimes\mathbf{B} \right)\boldsymbol{\cdot\Sigma} \right)^{-1}=\left( \begin{matrix} \text{se}\left( \hat{\beta}_{1} \right)^{2} & \cdots& \text{se}\left( \hat{\beta}_{1} \right)\cdot\text{se}\left( \hat{\beta}_{p} \right)\cdot\rho_{1p} \\ \vdots& \ddots& \vdots\\ \text{se}\left( \hat{\beta}_{p} \right)\cdot\text{se}\left( \hat{\beta}_{1} \right)\cdot\rho_{p1} & \cdots& \text{se}\left( \hat{\beta}_{p} \right)^{2} \end{matrix} \right)^{-1}=\frac{N_{Y}}{\hat{\sigma}_{Y}^{2}}\cdot\left( \begin{matrix} \frac{1}{\hat{\sigma}_{g,1}^{2}} & \cdots& \frac{\rho_{1p}}{\hat{\sigma}_{g,1}\cdot\hat{\sigma}_{g, p}} \\ \vdots& \ddots& \vdots\\ \frac{\rho_{p1}}{\hat{\sigma}_{g,1}\cdot\hat{\sigma}_{g, p}} & \cdots& \frac{1}{\hat{\sigma}_{g,p}^{2}} \end{matrix} \right)^{-1},$$

where $\text{B}$ denotes the vector of standard error of $\hat{\boldsymbol{\beta}}$’s and $\boldsymbol{\Sigma}$ denotes the square correlation coefficient structure of the SNPs. When all SNPs are independent, we can observe that $\boldsymbol{W}_{\text{MVIVW, correlated}}\boldsymbol{=}\boldsymbol{W}_{\text{MVIVW, independent}}$ since

$$\left( \begin{matrix} \text{se}\left( \hat{\beta}_{1} \right)^{2} & \cdots& \text{0} \\ \vdots& \ddots& \vdots\\ \text{0} & \cdots& \text{se}\left( \hat{\beta}_{p} \right)^{2} \end{matrix} \right)^{-1}=\left( \begin{matrix} \frac{1}{\text{se}\left( \hat{\beta}_{1} \right)^{2}} & \cdots& 0 \\ \vdots& \ddots& \vdots\\ 0 & \cdots& \frac{1}{\text{se}\left( \hat{\beta}_{p} \right)^{2}} \end{matrix} \right).$$

From the comparison, we could observe that the weight matrixes are different between hJAM and MVIVW MR.

#### Inverse-variance Weighted Mendelian Randomization

As described in Burgess et al. ^1^, the point estimate from inverse variance weighted Mendelian Randomization (IVW MR) can be expressed as

$$\hat{\pi}_{\mathrm{IVW}}= \frac{\sum_{k} \hat{\alpha}_{k}\hat{\beta}_{k}\cdot\text{s}\text{e}^{\text{-2}}(\hat{\beta}_{k})}{\sum_{k} \hat{\alpha}_{k}^{2}\cdot\text{s}\text{e}^{\text{-2}}\left( \hat{\beta}_{k} \right)}.$$

For SNP *k,* we have

$$se^{2}\left( \hat{\beta}_{k} \right)=\left( \boldsymbol{G}_{\boldsymbol{Yk}}^{\boldsymbol{'}}\boldsymbol{G}_{\boldsymbol{Yk}} \right)^{-1} \hat{\sigma}_{Y}^{2}=\frac{\hat{\sigma}_{Y}^{2}}{\sum\left( G_{i,Yk}-\bar{G}_{i,Yk} \right)^{2}}=\frac{\hat{\sigma}_{Y}^{2}}{g_{kk}^{2}}=\frac{\hat{\sigma}_{Y}^{2}}{{N_{Y}\hat{\sigma}}_{g,k}^{2}}.$$

Then,

$$\hat{\pi}_{\mathrm{IVW}}= \frac{\sum_{k} \hat{\alpha}_{k}\hat{\beta}_{k}\cdot\text{s}\text{e}^{\text{-2}}\left( \hat{\beta}_{k} \right)}{\sum_{k} \hat{\alpha}_{k}^{2}\cdot\text{s}\text{e}^{\text{-2}}\left( \hat{\beta}_{k} \right)}= \frac{\sum_{k} \hat{\alpha}_{k}\hat{\beta}_{k}\cdot{{\hat{\sigma}_{Y}^{-2} N}_{Y}\hat{\sigma}}_{g,k}^{2}}{\sum_{k} \hat{\alpha}_{k}^{2}\cdot{{\hat{\sigma}_{Y}^{-2} N}_{Y}\hat{\sigma}}_{g,k}^{2}}= \frac{\sum_{k} \hat{\alpha}_{k}\hat{\beta}_{k} \hat{\sigma}_{g,k}^{2}}{\sum_{k} \hat{\alpha}_{k}^{2} \hat{\sigma}_{g,k}^{2}}= \hat{\pi}_{\mathrm{hJAM}}.$$

and

$$\text{Var}\left( \hat{\pi}_{\mathrm{IVW}} \right)= \frac{1}{{\sum_{k} \hat{\alpha}}_{k}\cdot\frac{1}{\text{s}\text{e}^{\text{2}}\left( \hat{\beta}_{k} \right)}}= \frac{1}{{\sum_{k} \hat{\alpha}}_{k}\cdot\frac{{N_{Y}\hat{\sigma}}_{g,k}^{2}}{\hat{\sigma}_{Y}^{2}}}= \frac{\hat{\sigma}_{Y}^{2}}{{N_{Y}\cdot(\sum_{k} \hat{\alpha}}_{k}\cdot\hat{\sigma}_{g,k}^{2})}= \frac{\hat{\sigma}_{Y}^{2}}{N_{Y}\hat{\sigma}_{g}^{2}}.$$

Thus, the point estimate and its standard error that are estimated from IVW MR and *hJAM* are equivalent when $M=1$ and all instruments are independent.

#### Summary-PrediXcan

As described in ^3^, the point estimate from summary-PrediXcan (S-PrediXcan) can be expressed as

$$\hat{\pi}_{\text{Spred}}= \frac{\hat{\text{Cov}}\left( X, Y \right)}{\hat{\sigma}_{g}^{2}}= \frac{\hat{\text{Cov}}\left( \sum_{k} \alpha_{k}G_{k}, Y \right)}{\sigma_{g}^{2}}= \sum_{k} \frac{\hat{\text{Cov}}\left( \alpha_{k}G_{k}, Y \right)}{\hat{\sigma}_{g}^{2}}= \sum_{k} \frac{\hat{\alpha}_{k}\cdot\hat{\beta}_{k}\cdot\hat{\sigma}_{g,k}^{2}}{\hat{\sigma}_{g}^{2}}$$

$$=\frac{\sum_{k} \hat{\alpha}_{k}\hat{\beta}_{k} \hat{\sigma}_{g,k}^{2}}{\sum_{k} \hat{\alpha}_{k}^{2} \hat{\sigma}_{g,k}^{2}}$$

$$=\hat{\pi}_{\text{hJAM}}.$$

Note that summary-PrediXcan deals one gene per model. Thus, our point estimate is the same as summary-PrediXcan when $M=1$. Since PrediXcan uses elastic net to construct their $\boldsymbol{\alpha}$ vector, the association estimates in $\boldsymbol{\alpha}$ should be the joint SNP effects on the gene expression. From the properties of linear regression, we know that

$$\text{Var}\left( \hat{\pi}_{\text{Spred}} \right)=\frac{\hat{\sigma}_{\delta}^{2}}{N_{Y}\hat{\sigma}_{g}^{2}}=\frac{\hat{\sigma}_{Y}^{2}}{N_{Y}\hat{\sigma}_{g}^{2}}\left( 1-R_{g}^{2} \right)=\left( 1-R_{g}^{2} \right)\text{ Var}\left( \hat{\pi}_{\text{hJAM}} \right),$$

where $R_{g}^{2}$ denotes the heritability of gene expression $\boldsymbol{X}$. The Z test statistic of the significance of the association between the gene expression $\boldsymbol{X}$ and the trait $\boldsymbol{Y}$ can be expressed as:

$$Z_{\mathrm{Spred}}=\frac{\hat{\pi}_{\text{Spred}}}{\text{se}\left( \hat{\pi}_{\text{Spred}} \right)}=\sum_{k} \frac{\hat{\alpha}_{k}\hat{\beta}_{k} \hat{\sigma}_{g,k}^{2}}{\hat{\sigma}_{g}^{2}}\cdot\sqrt{\frac{N_{Y}\hat{\sigma}_{g}^{2}}{\hat{\sigma}_{Y}^{2}\left( 1-R_{g}^{2} \right)}}=\sum_{k} \frac{\hat{\alpha}_{k}\hat{\beta}_{k} \hat{\sigma}_{g,k}^{2}}{\hat{\sigma}_{g}}\cdot\sqrt{\frac{\left( 1-R_{g,k}^{2} \right)}{\text{se}\left( \hat{\beta}_{k} \right)^{2} \hat{\sigma}_{g,k}^{2}\left( 1-R_{g}^{2} \right)}}=\sum_{k} \frac{\hat{\alpha}_{k}\hat{\beta}_{k} \hat{\sigma}_{g,k}}{\hat{\sigma}_{g}\text{⋅ se}\left( \hat{\beta}_{k} \right)}\cdot\sqrt{\frac{\left( 1-R_{g,k}^{2} \right)}{\left( 1-R_{g}^{2} \right)}}\approx\frac{\sum_{k} \hat{\alpha}_{k}\hat{\beta}_{k} \hat{\sigma}_{g,k} \text{se}^{-1}\left( \hat{\beta}_{k} \right)}{\hat{\sigma}_{g}}=\frac{\sum_{k} \hat{\alpha}_{k} \hat{\beta}_{k}\hat{\sigma}_{g,k}^{2}}{\hat{\sigma}_{g}} \sqrt{\frac{N_{Y}}{\hat{\sigma}_{Y}^{2}}}$$

S-PrediXcan ignores the $\sqrt{\frac{\left( 1-R_{g,k}^{2} \right)}{\left( 1-R_{g}^{2} \right)}}$ term in the fourth line since they claim that this term does not affect their ability to detect the association based on their real data application and simulations. The $Z_{\mathrm{Spred}}$ statistic is same as the $Z_{\mathrm{hJAM}}$ statistic in our approach in the univariate case.

#### Summary-TWAS

The standard TWAS does not estimate $\hat{\pi}$ as other approaches do. It uses $Z_{TWAS}$ score to test the significance of the association between gene expression and the phenotypes. The weight matrix $W_{\text{TWAS}}$ in TWAS was compiled from the reference panel using summary statistics via ImpG-Summary algorithm ^7^. The $Z_{\text{TWAS}}$ statistic is computed as a linear combination of the standardized effect size, $\boldsymbol{Z}$*, of expression quantitative trait loci (eQTLs), i.e. for eQTL $k$,

$$Z_{k}^{*}=\frac{\hat{\beta}_{k}}{se(\hat{\beta}_{k})},$$

and the weight matrix $W_{\text{TWAS}}=\boldsymbol{\Sigma}_{\boldsymbol{G, X}}\boldsymbol{\Sigma}^{\boldsymbol{-1}}$, where $\boldsymbol{\Sigma}_{\boldsymbol{G,X}}$ denotes the covariance matrix between the genotype and the gene expression. The $Z_{\text{TWAS}}$ can be expressed as

$$Z_{\text{TWAS}}=\frac{{\boldsymbol{W}_{\text{TWAS}}\boldsymbol{Z}}^{*}}{\sqrt{\boldsymbol{W}_{\text{TWAS}}\boldsymbol{\cdot}\boldsymbol{\Sigma}\cdot\boldsymbol{W}_{\text{TWAS}}^{'}}}=\frac{\boldsymbol{\Sigma}_{\boldsymbol{G, X}}\boldsymbol{\Gamma}_{\mathbf{g}}^{\boldsymbol{-1}}\boldsymbol{\cdot}\boldsymbol{Z}^{\boldsymbol{*}}}{\sqrt{\boldsymbol{\Sigma}_{\boldsymbol{G, X}}\boldsymbol{\cdot}\boldsymbol{\Gamma}_{\mathbf{g}}^{\boldsymbol{-1}}\cdot\boldsymbol{\Sigma}_{\boldsymbol{G, X}}^{'}}}.$$

Here, for the eQTL $k$, assuming all the SNPs are independent, we have

$$\Sigma_{G_{k},X}=\text{Cov}\left( G_{k}, X \right)=\text{Cov}\left( G_{k}, \sum_{k} \alpha_{k}G_{k} \right)=\hat{\alpha}_{k}\hat{\sigma}_{g, k}^{2}.$$

Note that we have the ${(k, k)}^{th}$ diagonal element of $\boldsymbol{\Gamma}_{\boldsymbol{g}}^{\boldsymbol{-1}}$ to be $\frac{1}{\hat{\sigma}_{g,k}^{2}}$ under the independent assumption. Thus, we have

$$Z_{\text{TWAS}, independent}=\frac{\boldsymbol{\Sigma}_{\boldsymbol{G, X}}\boldsymbol{\Gamma}_{\mathbf{g}}^{\boldsymbol{-1}}\boldsymbol{\cdot}\boldsymbol{Z}^{\boldsymbol{*}}}{\sqrt{\boldsymbol{\Sigma}_{\boldsymbol{G, X}}\boldsymbol{\cdot}\boldsymbol{\Gamma}_{\mathbf{g}}^{\boldsymbol{-1}}\cdot\boldsymbol{\Sigma}_{\boldsymbol{G, X}}^{'}}}=\frac{\sum_{k} \hat{\alpha}_{k} \hat{\sigma}_{g,k}^{2}\cdot\frac{1}{\hat{\sigma}_{g,k}^{2}}\cdot\frac{\hat{\beta}_{k}}{\text{se}(\hat{\beta}_{k})}}{\sqrt{\sum_{k} \hat{\alpha}_{k}^{2} \hat{\sigma}_{g,k}^{2}}}=\frac{\sum_{k} \hat{\alpha}_{k} \hat{\beta}_{k}\text{se}^{-1}\left( \hat{\beta}_{k} \right)}{\hat{\sigma}_{g}}=\frac{\sum_{k} \hat{\alpha}_{k} \hat{\beta}_{k}\hat{\sigma}_{g,k}}{\hat{\sigma}_{g}} \sqrt{\frac{N_{Y}}{\hat{\sigma}_{Y}^{2}}},$$

which is slightly different from our estimate and S-PrediXcan: both hJAM and S-PrediXcan have an extra  $\hat{\sigma}_{g,k}$ in the numerator of the Z test score.
