## Supplementary figures and images for "A Hierarchical Approach Using Marginal Summary Statistics for Multiple Intermediates in a Mendelian Randomization or Transcriptome Analysis"

### Supplementary Figure 1

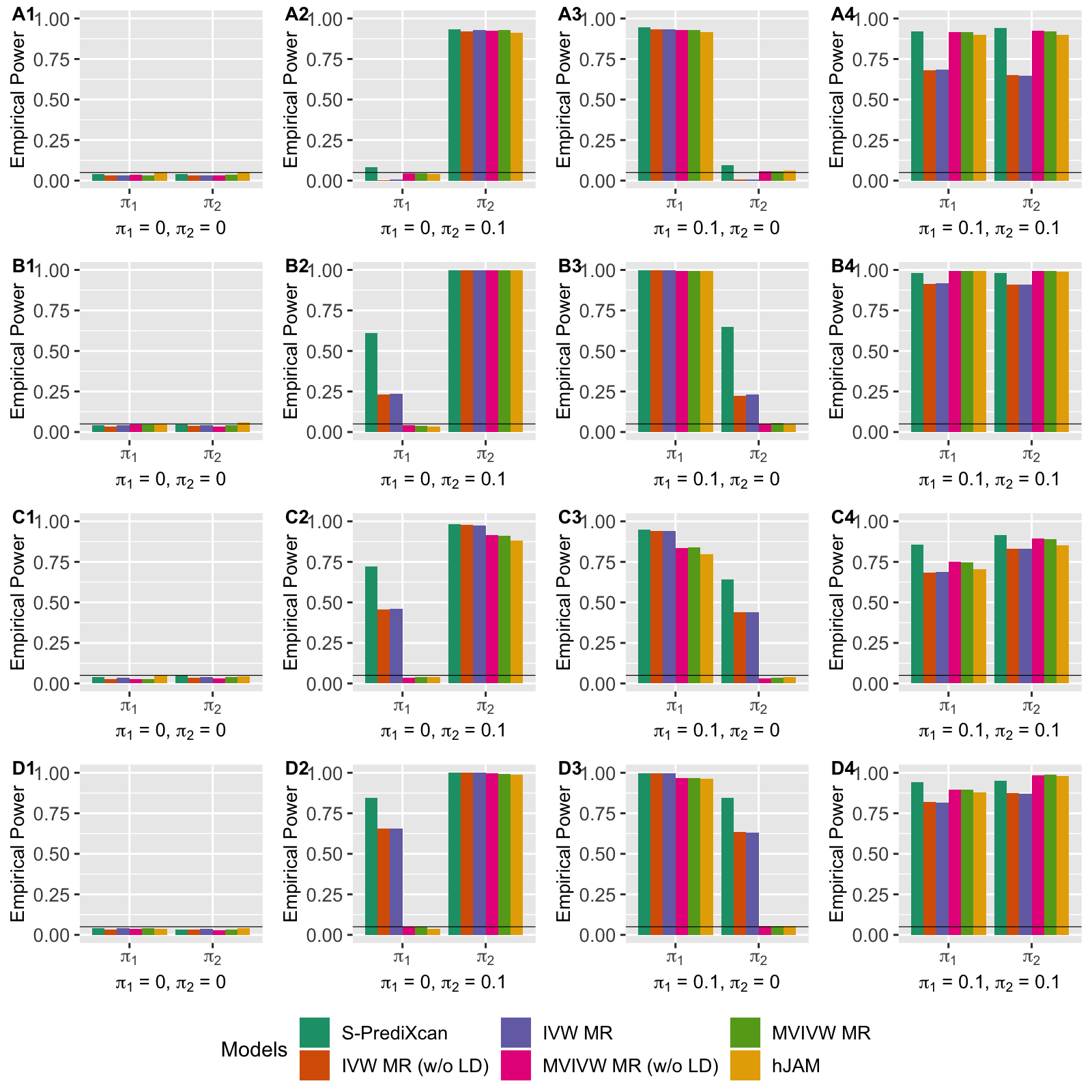

### Supplementary Figure 2

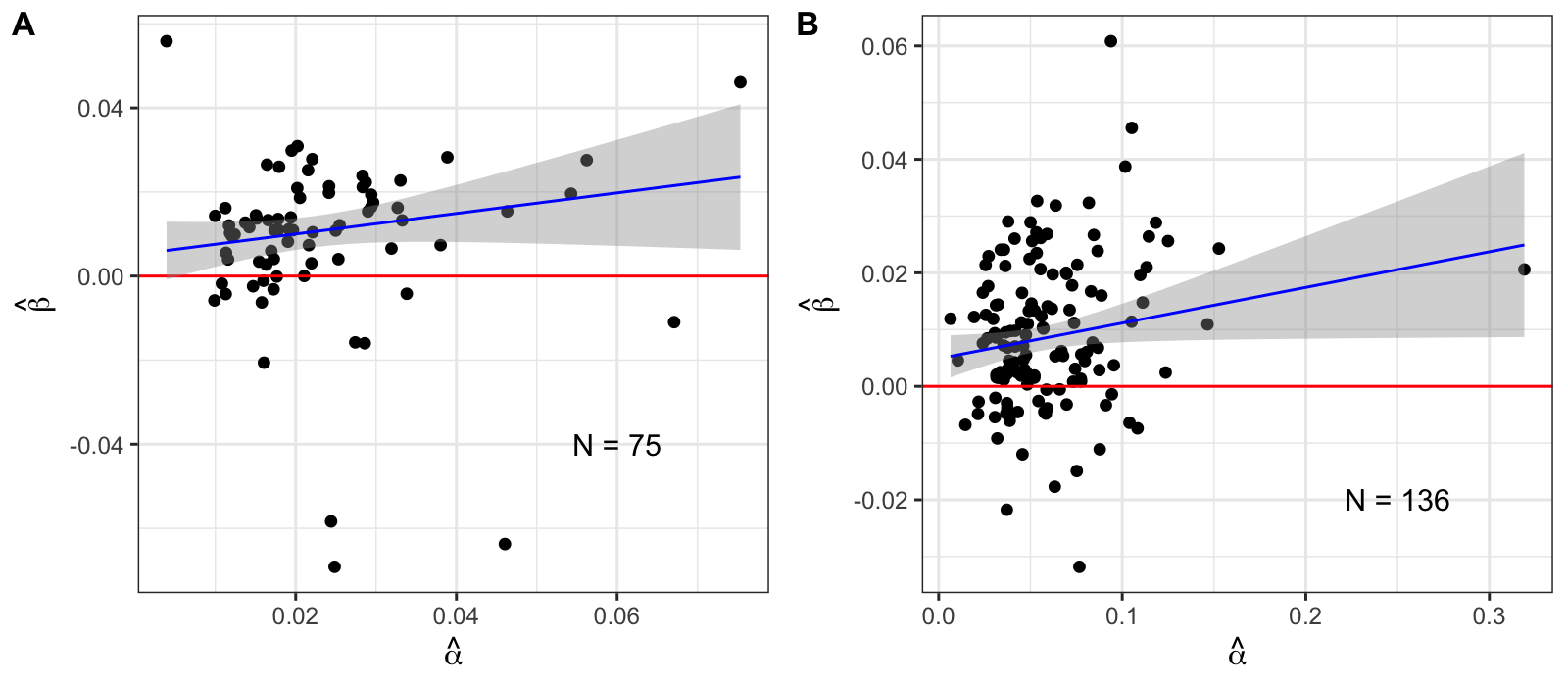

### Supplementary Figure 3

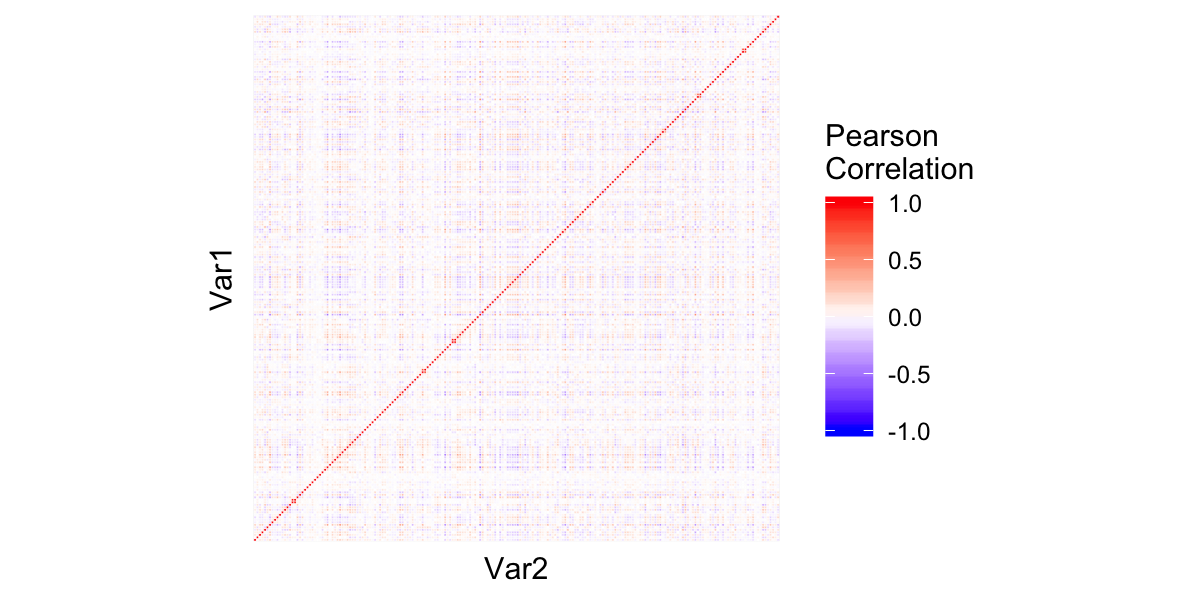

### Supplementary Figure 4

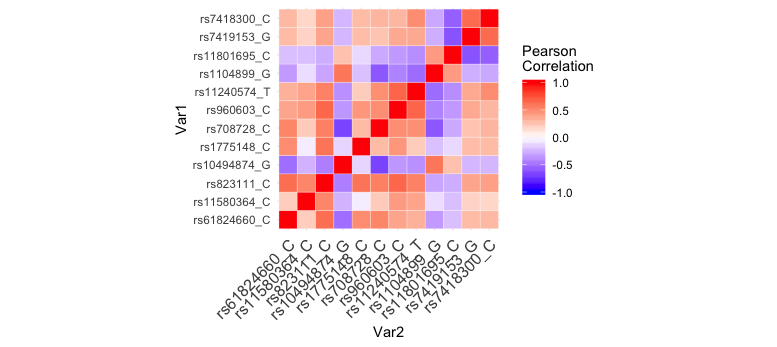
